# Unstructured Domains in Assembly Factors Promote the Hierarchy of Ribosome Maturation via Their Structural Plasticity

**DOI:** 10.1101/2023.11.23.568400

**Authors:** Ebba K. Blomqvist, Yoon-Mo Yang, Haina Huang, Katrin Karbstein

## Abstract

Ribosomes are assembled with the help of a large machinery of assembly factors (AFs), whose roles remain incompletely characterized. Recent structural studies of assembly intermediates have provided tremendous context for such studies and helped reveal novel roles, including as RNA chaperones. These structures have also revealed that a subset of the AFs have entirely non-globular structures, or large non-globular extensions, which extend across the assembling ribosomal subdomains, contacting many rRNA regions, ribosomal proteins (RPs) and other AFs. How these unusual structures could help promote assembly has remained unclear although it has been suggested that their potential for making multiple interactions helps constrain early assembly steps. By studying the roles of the AF Ltv1, an entirely non-globular protein, during late maturation steps of the 40S subunit’s head, we show here how the structural plasticity that derives from the non-globular structure is used to communicate maturation steps across the nascent subunit, thereby establishing the previously described hierarchy in head assembly. Our data indicate that while this structural plasticity enables integration of distinct folding and assembly steps, it also creates the potential for mutations that allow for bypass of these QC steps. These mutations are pathogenic in humans, further demonstrating the importance of proper 40S subunit assembly for protein homeostasis.

## Introduction

In all forms of life, ribosomes are responsible not just for producing the right amount of the correct protein, but also the detection of damaged mRNAs, as well as co-translational protein folding within the folding chamber and exit tunnel. All of these functions depend on the functional integrity of the ribosome pool, and efficient ribosome assembly, as reduced ribosome numbers or altered ribosome composition can affect both mRNA selection [1, 2], as well as its surveillance for damage [3].

In eukaryotes, ribosomes are assembled from their constituent 4 ribosomal RNAs (rRNAs) and 79 ribosomal proteins (RPs) with the help of ∼ 200 assembly factors (AFs), which are largely conserved from yeast to humans. These AFs integrate rRNA processing with binding of RPs, chaperone rRNA folding, allow for regulation, prevent premature translation initiation, and facilitate quality control [4–8].

Ribosome assembly overall is a hierarchical process, leading to the formation of a series of assembly intermediates that can be ordered based on their composition. This hierarchy is not limited to the association of proteins, both RPs and AFs, but also to their dissociation. One advantage of hierarchical assembly is that it enables the establishment of quality control checkpoints. If there are many routes to a mature subunit, and are many valid intermediates, progression of assembly intermediates lacking a critical piece is harder to prevent, as they could be similar or identical to on-pathway intermediates from other routes. In contrast, if assembly is very hierarchical, it is simpler to establish a pass-fail system, and sieve out intermediates that fail to complete any given step. It is also likely that the hierarchy established by AFs could support hierarchical RP assembly beyond the waves that were observed in *in vitro* systems that lack those factors [9].

A subset of AFs lack globular domains, and instead are made up of a series of helices and small beta sheets, which thread throughout assembly intermediates, connecting distant parts. These include Utp11, Utp14, Nop14, Mpp10, as well as Nog1, Nog2, Ebp1 and others for the small and large subunit, respectively. Many additional AFs (and RPs) have long non-globular extensions to their globular structures. If and how these unusual structures are used to connect distinct sites and facilitate hierarchy and quality control is not known, although it has been speculated that they might allow for the compaction [6] or integration of processing events [10].

One of these non-globular AFs is Ltv1 (**Figure 1A&B**). After binding in the nucleolus, Ltv1 accompanies nascent 40S subunits into the cytoplasm [11], where it is one of seven stably bound factors present in late assembly intermediates. Indeed, it is the Hrr25/CK1δ-mediated phosphorylation and subsequent release of Ltv1 that commits the late pre-40S intermediate into a translation-like cycle [12–16], which integrates maturation with quality control to ensure that only fully and correctly matured ribosomes enter the translating pool [7, 16–19]. Specifically, binding of the RPs Rps3, Rps15, Rps20, Rps29 and Rps31, and folding of the rRNA is integrated with phosphorylation of two AFs, Ltv1 and Rio2, as well as the ordered dissociation of Ltv1, Enp1 and Rio2 (**Figure 1C**, [9, 16, 20]). This hierarchy underlies a crucial quality control checkpoint, which ensures that nascent ribosomes are able to correctly identify the start codon during translation initiation [16]. Importantly, disrupting the hierarchy allows for release of misassembled ribosomes into the translating pool [9, 16, 21], leading to defects in start codon selection and demonstrating its importance for ribosome integrity and cellular protein homeostasis.

**Figure 1:**
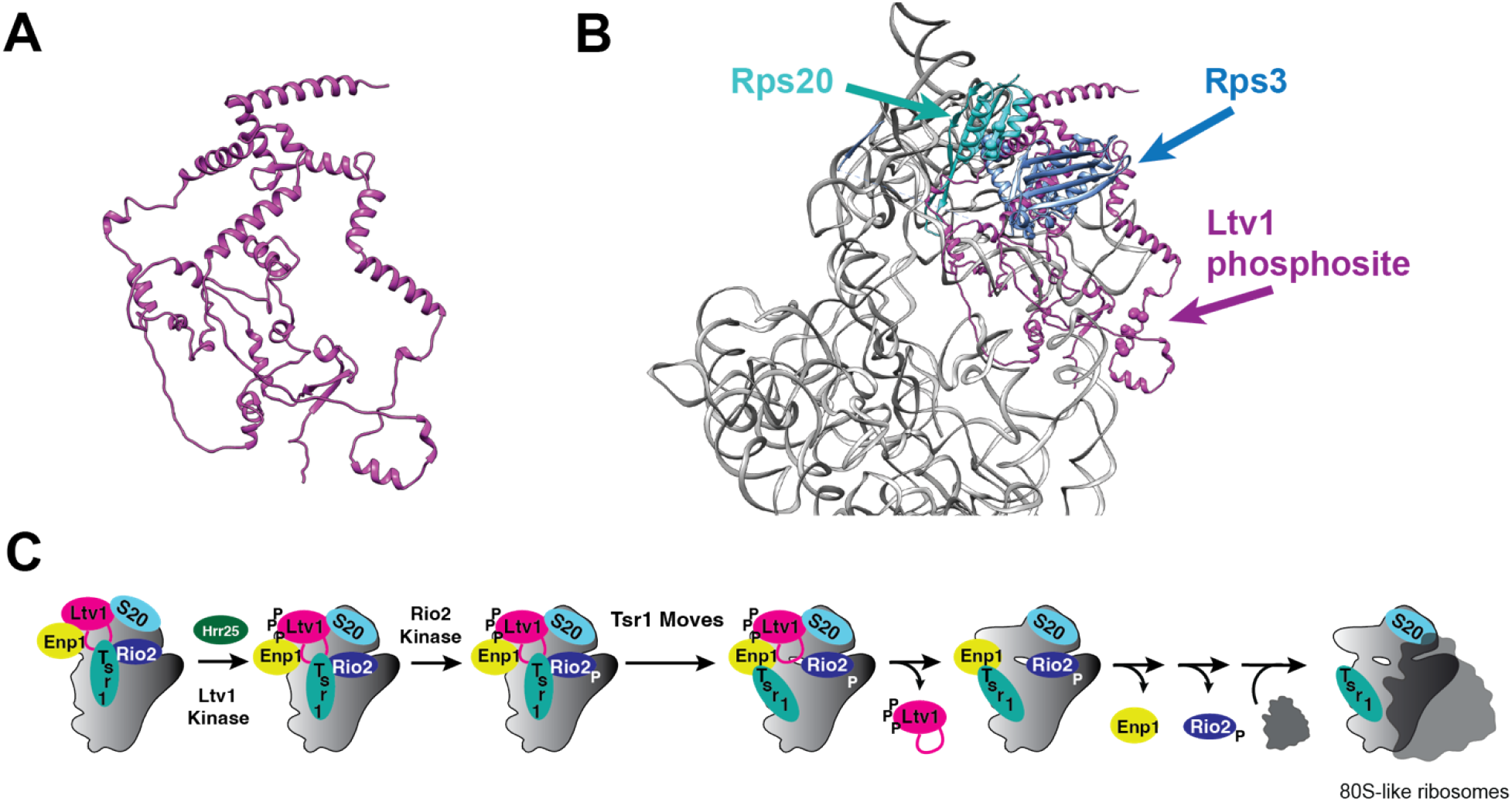
Ltv1 is involved in the hierarchical cascade of events that link quality control of head formation to the formation of 80S-like ribosomes. (A) Alpha-fold predicted structure of Ltv1 [34, 35]. (B) Predicted placement of Ltv1 on pre-40S ribosomes. Adapted from [9]. Residues in Rps3 and Rps20 that interact in mature 40S are indicated in spacefill. The residues in Ltv1 that are phosphorylated by Hrr25 for their release [13, 15] are also indicated in spacefill. (C) Steps that convert stable cytoplasmic pre-40S ribosomes into 80S-like ribosomes. Adapted from [9, 16].

Despite the fact that this hierarchy in assembly has been characterized in detail, and despite its importance for quality control, it remains unclear how the individual events are communicated across the assembling subunit to enable the next step, thereby establishing hierarchy. Here we use a collection of genetic and biochemical analyses to dissect how the cascade of late assembly steps that commits nascent subunits to maturation is initiated. Our data show that via its flexible, multi-point interactions with the small ribosomal subunit Ltv1 communicates assembly steps across the subunit, thereby enabling hierarchy in small subunit head assembly. Bypass of individual steps is observed in cancer cells. These observations may be of general relevance for non-globular AFs.

## Results

We have recently shown that Ltv1 initially blocks the premature formation of the contact between residues K7 and K10 of Rps3 and D113 and D115 of Rps20 (**Figure 1B**, [9]). However, we and others have also shown that these contacts are required for 40S maturation [13, 16], specifically for phosphorylation of Ltv1 (**Figure 2A**, [16]), which initiates a cascade of events that ultimately leads to the formation of 80S-like ribosomes (**Figure 1C**, [15]). 80S-like ribosomes are critical intermediates in 40S assembly, which couple maturation to the successful completion of translation-like steps as part of a quality control (QC) mechanism [16, 17, 22]. Thus, we wondered how Ltv1 and/or Rps3/Rps20 were repositioned during 40S maturation to establish these contacts and initiate the final maturation cascade.

**Figure 2:**
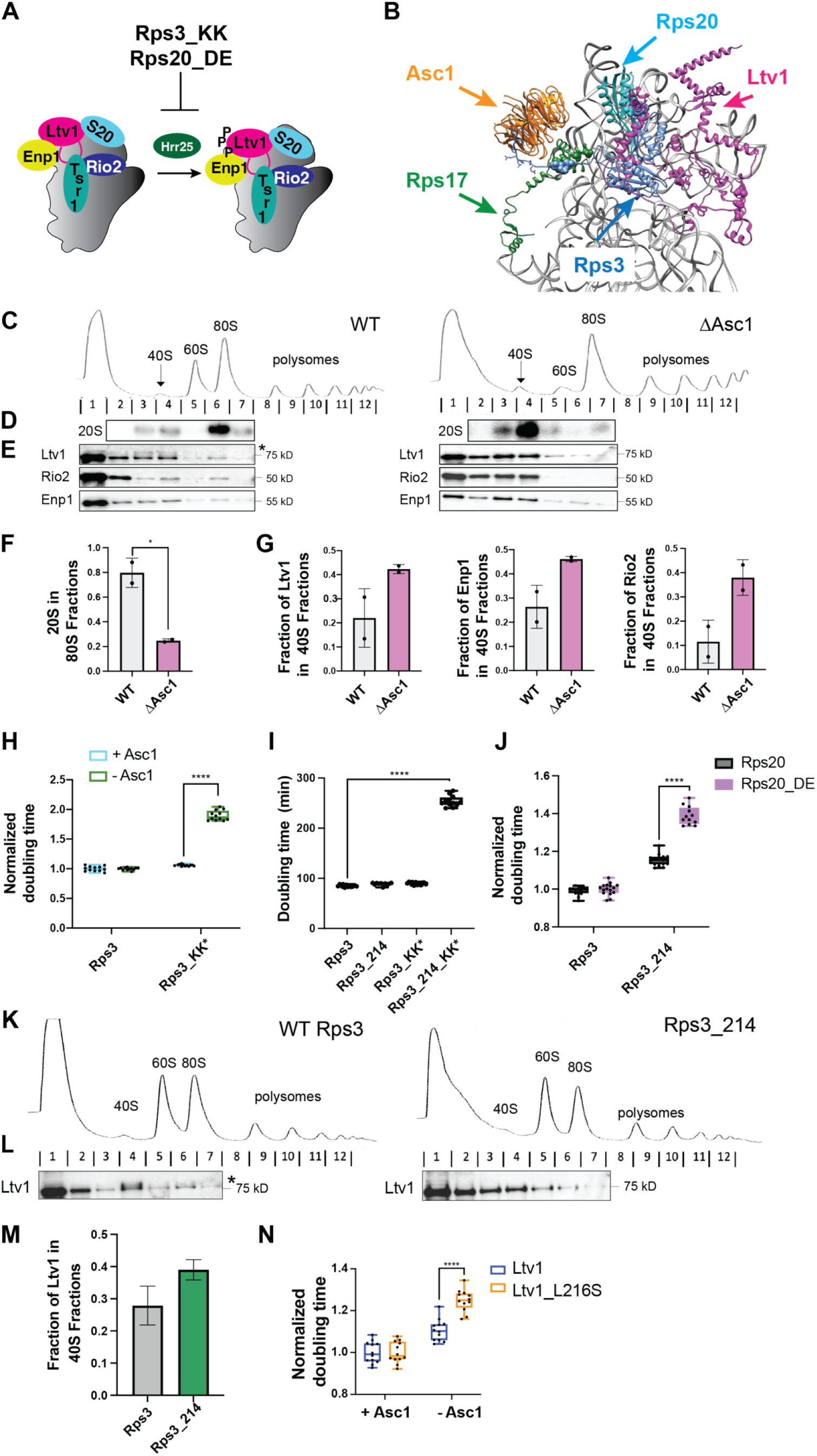
Asc1 is required for Ltv1 phosphorylation and release. (A) Excerpt of the cascade in Figure 1C that highlights the Ltv1 phosphorylation step, which is blocked by the Rps3_KK and Rps20_DE mutations. (B) Detail of pre-40S ribosomes (pdb ID 6FAI), with Rps3 and Asc1 in the mature positions (from pdb ID 3JAM), and Ltv1 placed as in [9]. The Rps3_IEP and Rps17_LRYTQ residues that interact in mature ribosomes are highlighted in blue and green spacefill, respectively. The Rps3_KK and Rps20_DE residues that interact in mature ribosomes are highlighted in blue and cyan spacefill, respectively. (C) Absorbance profiles at 260 nm of 10-50% sucrose gradients from Fap7 depleted cells either without or with the Asc1 deletion. The U24 snoRNA in the Asc1 intron was provided on a plasmid. (D) Northern and (E) Western blots of the gradient fractions in panel C. The position of phosphorylated Ltv1 is indicated with the asterisk. (F) Quantification of the Northern blots in panel D and a biological replicate. (G) Quantification of the Western blots in panel E and a biological replicate. Data are shown as a mean with standard deviation. (H) Normalized (to WT Rps3) doubling times from yeast expressing wt Rps3, or Rps3_KK in the presence or absence of Asc1. Significance was tested using a two-way ANOVA. ****, P<0.0001. (I) Doubling times from yeast expressing wt Rps3, the C-terminal truncation at residue 214 (Rps3_C214), Rps3_KK or Rps3_C214_KK. Rps3_C214 disrupts Asc1 binding [25]. Significance was tested using a two-way ANOVA. ****, P<0.0001. (J) Normalized (to WT Rps3) doubling times from yeast expressing wt Rps3, or Rps3_C214 in the presence of wt Rps20 or Rps20_DE. Significance was tested using a two-way ANOVA. ****, P<0.0001. (K) Absorbance profiles at 260 nm of 10-50% sucrose gradients from Fap7 depleted cells expressing either wt Rps3 or Rps3_C214. (L) Western blots for Ltv1 of the gradient fractions in panel K. The position of phosphorylated Ltv1 is indicated with the asterisk. (M) Quantification of the Western blots in panel L and a biological replicate. Data are shown as a mean with standard deviation. (N) Normalized (to +Asc1) doubling times from yeast expressing wt Ltv1, or Ltv1_216S and either containing or lacking Asc1. Significance was tested using a two-way ANOVA. ****, P<0.0001.

### Asc1 repositions Rps3 to establish the contact between Rps3 and Rps20

In mature 40S subunits Rps3 binds two other ribosomal proteins (RPs), Rps17 and Asc1 (**Figure 2B**). Moreover, Asc1 is recruited sometime in the transition to 80S-like ribosomes, as it is not found in pre-40S ribosomes prior to Ltv1 phosphorylation [15, 23], but appears fully bound in 80S-like ribosomes [24]. Furthermore, deletion of Ltv1 leads to partial loss of Asc1 from ribosomes, likely mediated by Rps3 [25]. Given these observations, we asked whether Asc1 binding to pre-40S ribosomes could reposition Ltv1, leading to its phosphorylation. To test this hypothesis, we created a yeast strain in which Asc1 was deleted and supplemented this strain with a plasmid encoding U24 snoRNA, which is normally encoded in the intron of the Asc1 gene. We first wanted to test whether deletion of Asc1 impaired the formation of 80S-like ribosomes. Thus, we modified this yeast strain further by introducing a galactose-inducible promoter in front of the Fap7 gene, to allow for depletion of Fap7, which leads to the accumulation of 80S-like ribosomes [16, 17, 22], unless their formation is otherwise blocked. Thus, by comparing the sedimentation of pre-40S ribosomes, which contain 20S pre-rRNA, the direct precursor to 18S rRNA, in Fap7 depleted strains with and without Asc1, we can determine whether Asc1 is required for the formation of 80S-like ribosomes, as previously described [16, 17, 22].

Indeed, the absence of Asc1 shifts the equilibrium between 80S-like and 40S ribosomes from 80S-like ribosomes to earlier 40S ribosomes (**Figure 2C&F**), demonstrating that Asc1 is required for the efficient formation of 80S-like ribosomes. Next, we wanted to narrow the timing of Asc1 binding during the series of events that starts with Ltv1 phosphorylation and ends in the formation of 80S-like ribosomes. We have previously shown that Ltv1 phosphorylation by Hrr25 initiates a cascade of events that leads to the ordered release of Ltv1, Enp1 and Rio2 and the formation of 80S-like ribosomes (**Figure 1C**, [16]). We therefore measured the accumulation of these three assembly factors (AFs) in pre-40S ribosomes utilizing western blots over the gradients in **Figure 2C**. This analysis demonstrates that Asc1 depletion leads to the accumulation of Ltv1, Enp1 and Rio2 on pre-ribosomes (**Figure 2E&G)**, indicating that Asc1 acts upstream of the first of these events, the release of Ltv1. Finally, in the absence of Asc1, no Ltv1 phosphorylation can be detected (**Figure 2E**), demonstrating that Asc1 is required for phosphorylation of Ltv1. This observation supports the model discussed above that Asc1 moves Ltv1, and/or Rps3/Rps20 to help establish the contact between Rps3 and Rps20.

To further support this model, we tested for genetic interactions between Asc1 deletion and mutation of Rps3. Indeed, mutation of Rps3 K7 and K10 to alanine (Rps3_KK*), does not demonstrate a growth defect in the presence of Asc1, but shows a significant synthetic growth defect with Asc1 deletion (**Figure 2H**), as expected if Asc1 moves Ltv1 to establish the contact between the Rps3_KK residues and Rps20.

We have previously shown that truncation of Rps3 at residue 214 (Rps3_214) leads to loss of Asc1 from ribosomes [25]. We therefore confirmed that truncation of Rps3 at residue 214 together with Rps3_KK demonstrates the same synthetic negative effect that was observed from the combination of Rps3_KK and Asc1 deletion (**Figure 2I**). Having validated that as expected, we next used the Rps3_214 truncation and combined it with Rps20_DE, the residues that bind Rps3_KK in mature ribosomes. As expected Rps20_DE is also synthetically sick with truncation of Rps3 (**Figure 2J**), which disables Asc1 binding [25], thereby supporting the model that Asc1 helps establish the contact between Rps3 and Rps20.

To further investigate the role of Asc1 in enabling the contact between Rps3 and Rps20 further, we assessed if deletion of the Asc1 binding site in Rps3 has the same effect as deletion of Asc1. We therefore tested for Ltv1 phosphorylation and release in yeast strains expressing Rps3_214. As described above, we utilized a yeast strain in which 80S-like ribosomes accumulate via the depletion of Fap7 and combined this with a galactose-inducible/glucose-repressible Rps3 strain, to produce either wild type or Rps3_214 from plasmids, and then utilized sucrose gradients to fractionate the lysates from these strains after Fap7 depletion (**Figure 2K**). As expected, Ltv1 phosphorylation is lost in the Rps3_214 strain (**Figure 2L**), which lacks Asc1, leading to the expected accumulation of Ltv1 in pre-40S ribosomes (**Figure 2L-M**).

Finally, Asc1 deletion is synthetically sick with Ltv1_L216S (**Figure 2N**), which also impairs Ltv1 phosphorylation and is synthetically sick with Rps3_KK* and Rps20_DE [9]. Our previous analysis has shown that Ltv1 is mispositioned by this mutation, in particular allowing for the premature formation of the contact between Rps3 and Rps20. Thus, the synthetic negative interactions between Asc1 deletion and Ltv1_L216S, and Asc1 deletion and Rps3_KK* both suggest that Asc1 acts on Rps3, consistent with its interaction with Rps3 [25].

Together, these experiments demonstrate that Asc1 is required for phosphorylation of Ltv1 and strongly suggest that it functions by repositioning Rps3, thereby establishing the contact between Rps3 and Rps20.

### Rps17 aids Asc1 in repositioning of Rps3

We have previously shown that the Rps17_LRYTQ mutation (L16A,R19E,Y20A,T39A,Q42A, **Figure 2A**), which impairs the contact between Rps3 and Rps17, is epistatic with deletion of Ltv1 [25]. The same is true for the corresponding mutation in Rps3, Rps3_IEP (I208A, E210R and P211A), which blocks Rps17 binding, [25]. In addition, other previous work indicated that Rps17 is repositioned late in assembly, around the time when 80S-like ribosomes are formed [26]. We therefore investigated whether Rps17 is also linked to repositioning of Rps3.

We produced yeast strains where Fap7 could be depleted by growth in dextrose, and expressing wild type Rps17, or the Rps17 mutant defective for binding to Rps3, Rps17_LRYTQ. We then used these to test for the formation of 80S-like ribosomes as described above. As observed for depletion of Asc1, disrupting the Rps3_Rps17 contact shifts the equilibrium from 80S-like to pre-40S intermediates, indicating that the mutation blocks the formation of 80S-like ribosomes (**Figure 3A, B, D**). In addition, in yeast expressing Rps17_LRYTQ, Ltv1, Enp1 and Rio2 accumulate on pre-ribosomes, and Ltv1 phosphorylation is absent (**Figure 3C&E**).

**Figure 3:**
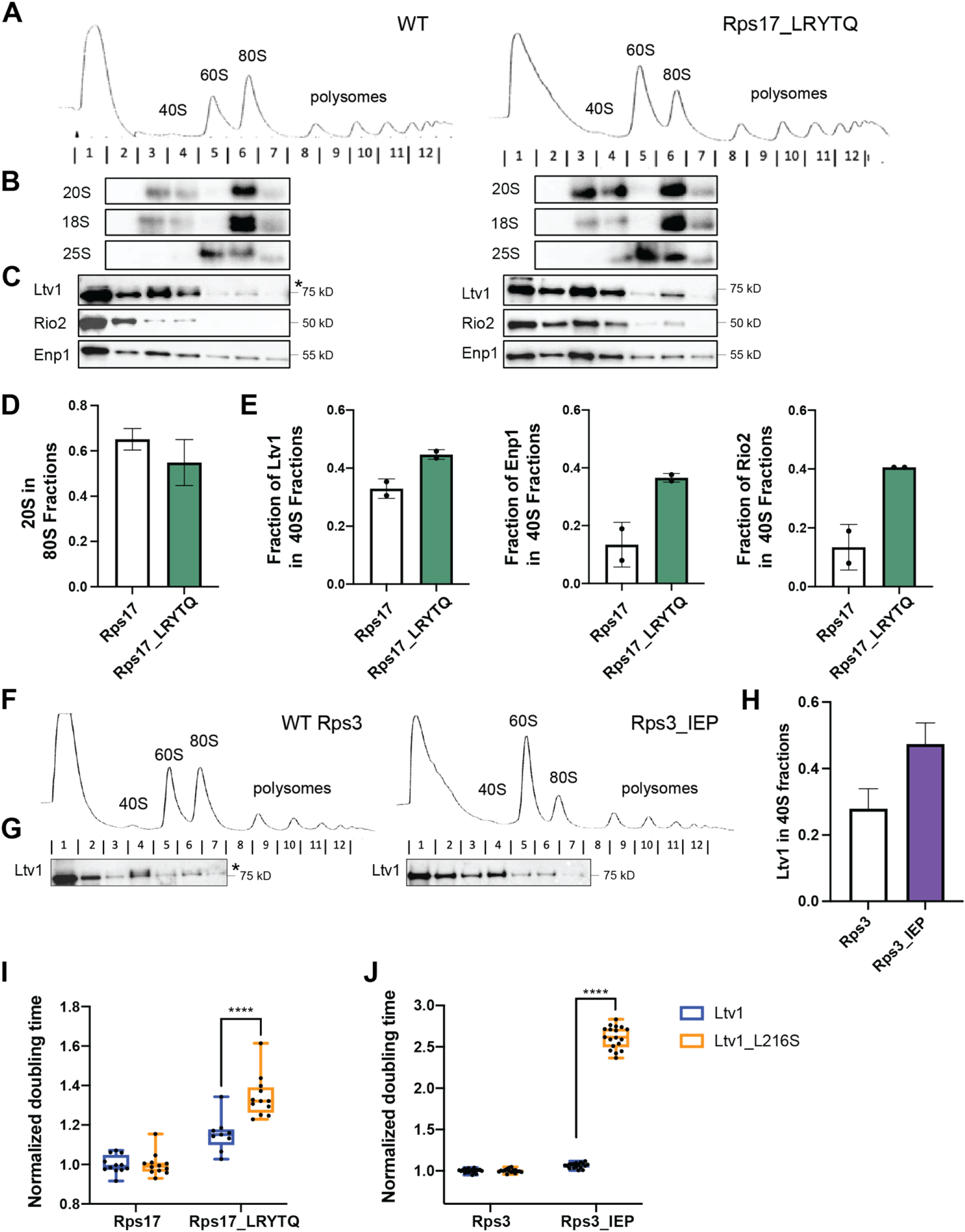
Rps17 is required for Ltv1 phosphorylation and release. (A) Absorbance profiles at 260 nm of 10-50% sucrose gradients from Fap7 depleted cells expressing either wt Rps17 or Rps17_LRYTQ, which does not bind Rps3 (Figure 2B). (B) Northern and (C) Western blots of the gradient fractions in panel A. The position of phosphorylated Ltv1 is indicated with the asterisk. (D) Quantification of the Northern blots in panel B and a biological replicate. (E) Quantification of the Western blots in panel C and a biological replicate. Data are shown as a mean with standard deviation. (F) Absorbance profiles at 260 nm of 10-50% sucrose gradients from Fap7 depleted cells expressing either wt Rps3 or Rps3_IEP, which does not bind Rps17 (Figure 2B). (G) Western blots of the gradient fractions in panel E. The position of phosphorylated Ltv1 is indicated with the asterisk. (H) Quantification of the Western blots in panel F and a biological replicate. Data are shown as a mean with standard deviation. (I) Normalized (to wt Rps17) doubling times from yeast expressing wt Ltv1, or Ltv1_216S and wt Rps17 or Rps17_LRYTQ. Significance was tested using a two-way ANOVA. ****, P<0.0001. (J) Normalized (to wt Rps17) doubling times from yeast expressing wt Ltv1, or Ltv1_216S and wt Rps17 or Rps17_LRYTQ. Significance was tested using a two-way ANOVA. ****, P<0.0001.

Similarly, when we construct yeast strains expressing wild type Rps3 or Rps3_IEP, which mutates the contact for Rps17_LRYTQ, and carry out sucrose gradient analysis in the absence of Fap7, we also find that Ltv1 phosphorylation is lost, and Ltv1 accumulates on pre-ribosomes (**Figure 3F-H**).

Finally, like the deletion of Asc1, Rps17_LRYTQ is synthetically sick with Ltv1_L216S (**Figure 3I**). Moreover, the corresponding mutation in Rps3, Rps3_IEP, also demonstrates synthetically sick phenotypes with Ltv1_216 (**Figure 3J**). Thus, the contact between Rps17 and Rps3 is required for phosphorylation of Ltv1, akin to the contact between Asc1 and Rps3.

Together, these data demonstrate that Asc1 and Rps17, via their interactions with Rps3, are required for phosphorylation of Ltv1, likely by repositioning Rps3.

### Ltv1 links changes in Rps3 and Rps20 to the phosphosite

Our previous data demonstrate that Ltv1 blocks the premature formation of the contact between Rps3 and Rps20 [9]. Above, we have shown that binding of Asc1 and Rps17 to Rps3 are required to establish this contact, likely by repositioning at least Rps3. Because we had also shown that formation of this contact was required for Ltv1 phosphorylation [16], and 40S maturation [13], we next asked how this conformational rearrangement was communicated to the phosphorylation site in Ltv1 (**Figure 4A**). In principle, this could occur through changes in the pre-40S structure, induced by Rps3 repositioning. In addition, because Rps3 interacts with Ltv1 via at least two independent binding sites [13, 27], it was also possible that Ltv1 alters its own structure, thereby communicating this contact directly to its phosphorylation site, which is distant in both primary sequence and tertiary structure, with Rps3/Rps20 being positioned on the back of the head, and the phosphorylation site being located near Rps12 on the side of the beak (**Figure 1B**).

**Figure 4:**
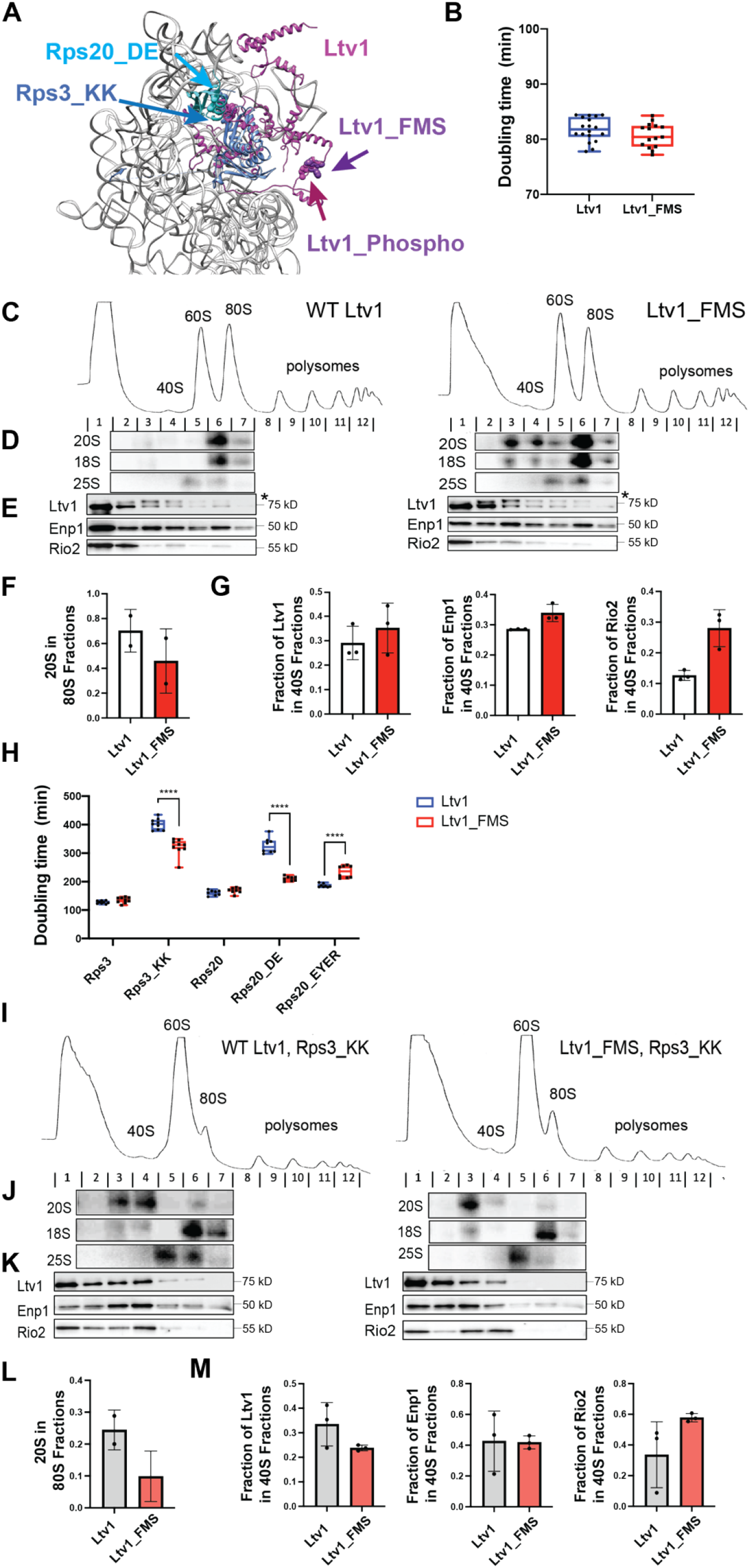
Ltv1 communicates the contact between Rps3 and Rps20 to its phosphorylation site. (A) Detail of pre-40S ribosomes (pdb ID 6FAI) with Ltv1 placed as in [9]. The Rps3_KK and Rps20_DE residues that interact in mature ribosomes are highlighted in blue and cyan spacefill, respectively. The Ltv1_FMS residues and the pshospho-site are adjacent and in purple and magenta spacefill, respectively. (B) Doubling times from yeast expressing wt Ltv1 or Ltv1_FMS. (C) Absorbance profiles at 260 nm of 10-50% sucrose gradients from Fap7 depleted cells expressing either wt Ltv1 or Ltv1_FMS. (D) Northern and (E) Western blots of the gradient fractions in panel C. The position of phosphorylated Ltv1 is indicated with the asterisk. (F) Quantification of the Northern blots in panel D and a biological replicate. (G) Quantification of the Western blots in panel E and a biological replicate. Data are shown as a mean with standard deviation. (H) Doubling times from yeast expressing wt Ltv1 or Ltv1_FMS and either wt Rps3, or Rps3_KK or wt Rps20 or Rps20_DE, or Rps20_EYER. Significance was tested using a two-way ANOVA. ****, P<0.0001. (I) Absorbance profiles at 260 nm of 10-50% sucrose gradients from Fap7 depleted cells expressing either wt Ltv1 or Ltv1_FMS and Rps3_KK. (J) Northern and (K) Western blots of the gradient fractions in panel I. The position of phosphorylated Ltv1 is indicated with the asterisk. (L) Quantification of the Northern blots in panel I and a biological replicate. Data are shown as a mean with standard deviation. (M) Quantification of the Western blots in panel J and a biological replicate. Data are shown as a mean with standard deviation.

To test if Ltv1 itself was involved in communicating the formation of the Rps3/Rps20 contact to its phosphorylation site, we made mutations around the Ltv1 phosphorylation site, mutating F341, M343 and S344 to alanine (the phosphorylated residues are S336, 339 and 342), to produce Ltv1_FMS (**Figure 4A**). Ltv1_FMS has essentially no growth phenotype by itself (**Figure 4B)**, although sucrose gradient analysis demonstrates a small defect in the formation of 80S-like ribosomes (**Figure 4C&F)** and a small accumulation of Rio2 in pre-40S subunits, reflecting a defect in Rio2 release (**Figure 4E, G)**.

Next, we tested for genetic interactions between Ltv1_FMS and Rps3_KK and Rps20_DE. To produce a growth defect, we utilized the Rps3_K7E, K10D mutation (not the corresponding alanine mutants, Rps3_KK*) Notably, Ltv1_FMS partially rescues the growth defects of both Rps3_KK and Rps20_DE (**Figure 4H)**, but not of Rps20_EYER, which impairs the recruitment of Rps20 via mutation of its interface with Rps29 [9, 16].

To test if Ltv1_FMS also rescued the formation of 80S-like ribosomes and the release of Ltv1, Enp1 and Rio2 in Rps3_KK cells, we used sucrose gradient analysis as described above (**Figure 4I-M)**. This analysis demonstrates that Ltv1_FMS indeed rescues the accumulation of Ltv1 and Enp1 in pre-40S ribosomes. In contrast, Rio2 remains accumulated in pre-40S ribosomes (**Figure 4K&M**), explaining while the formation of 80S-like ribosomes remains impaired (**Figure 4J&L**). Thus, changing the structure of Ltv1 around the phosphorylation site rescues the detrimental effects from blocking the Rps3-Rps20 contact, indicating that conformational changes within the ribosome are not required to communicate the formation of the Rps3-Rps20 contact to the Ltv1 phosphorylation site, but that rather Ltv1 mediates the communication. Note that Rio2 accumulating on pre-40S ribosomes also suggests that either communication to the C-terminal tail of Rps15, where all other mutations that block Rio2 release are located [16], is broken in this mutant or that Rio2 fails to be phosphorylated.

### Ltv1 communicates its phosphorylation to changes in l31 to enable its release

We have previously shown that phosphorylation of Ltv1 initiates a hierarchical set of events, which include the release of Ltv1, Enp1 and Rio2, and culminate in the formation of 80S-like ribosomes. However, our previous data has also shown that phosphorylation and release of Ltv1 were temporally separated, and also required the phosphorylation of Rio2 [16] and an additional step, which we have since identified as repositioning of Tsr1 from the decoding helix to the beak ([9], **Figure 1C & 5A**). How these events are communicated to affect the next step remained unknown.

We have previously shown that Rio2 bound directly to a late-folding RNA element in the 40S head, the loop of helix 31 (l31) [20]. Moreover, we have recently shown that l31 also binds directly to Ltv1, an interaction that is perturbed by mutation of the Ltv1 residues Y82, D83, and Y84 to VAV to produce Ltv1_YDY (**Figure 5B**, [9]). We therefore hypothesized that l31 established communication between Rio2 and Ltv1 phosphorylation and release.

**Figure 5:**
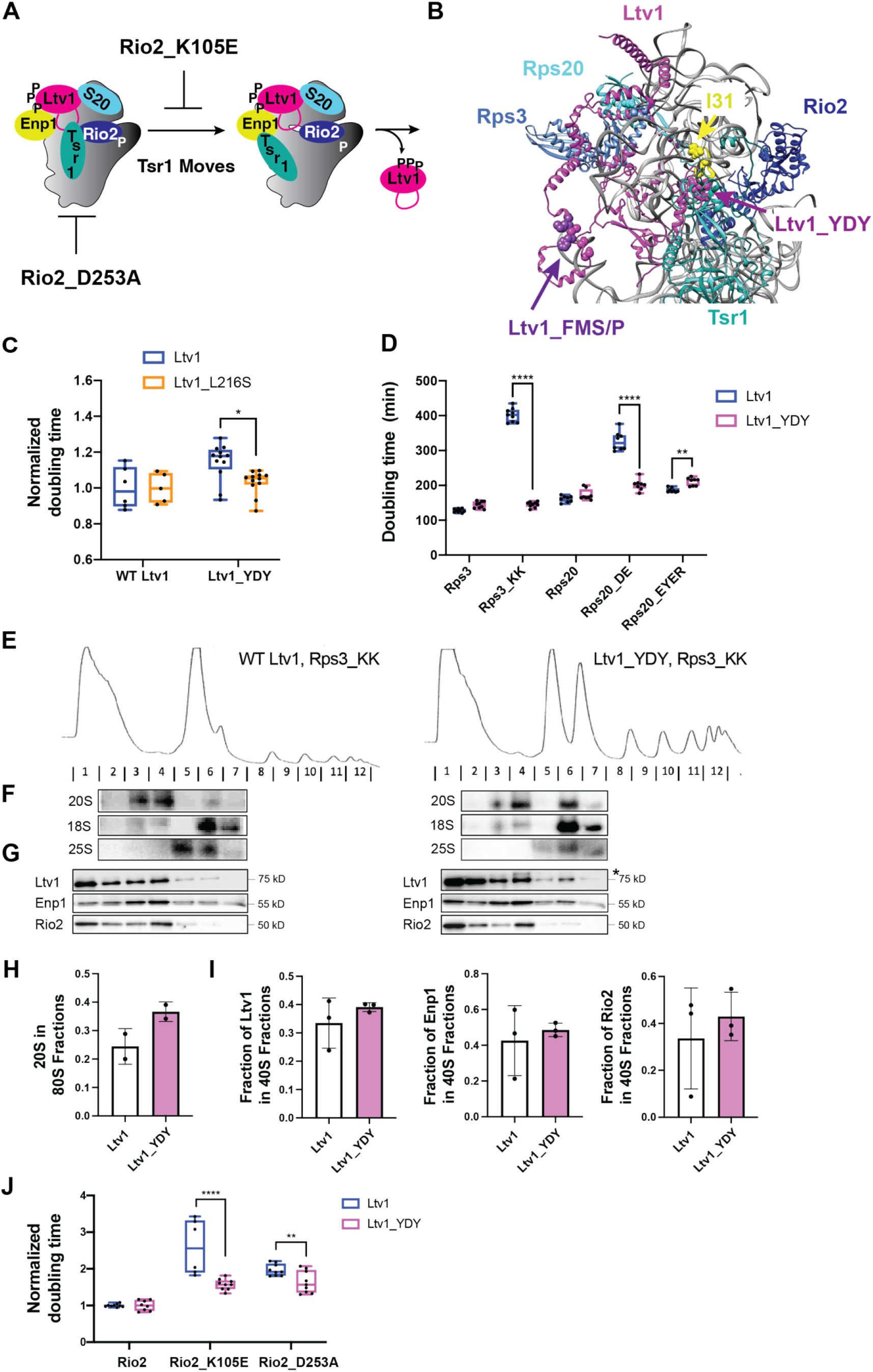
Ltv1 communicates it phosphorylation to l31. (A) Excerpt of the cascade in Figure 1C that highlights the Tsr1 movement, which is required for Ltv1 release and blocked by the Rio2_K105E mutation [9]. Rio2_D253A is upstream and blocks Rio2 phosphorylation. (B) Detail of pre-40S ribosomes (pdb ID 6FAI), and Ltv1 placed as in [9]. The Rps3_KK and Rps20_DE residues that interact in mature ribosomes are highlighted in blue and cyan spacefill, respectively. The Ltv1_FMS residues and the pshospho-site are adjacent and in purple and magenta spacefill, respectively. Tsr1 is in cyan, Rio2 in blue. The l31 loop in 18S rRNA is disordered and its first and last residues are shown in yellow spacefill. Ltv1_YDY is in purple spacefill. (C) Normalized (to wt Ltv1) doubling times from yeast expressing wt Ltv1, Ltv1_L216S, Ltv1_YDY, or Ltv1_YD_L216S. Significance was tested using a two-way ANOVA. *, P<0.05. (D) Doubling times from yeast expressing wt Ltv1, or Ltv1_YDY and wt Rps3, Rps3_KK, wt Rps20, or Rps20_DE or Rps20_EYER. Significance was tested using a two-way ANOVA. **, P<0.01; ****, P<0.0001. (E) Absorbance profiles at 260 nm of 10-50% sucrose gradients from Fap7 depleted cells expressing Rps3_KK and either wt Ltv1, or Ltv1_YDY. (F) Northern and (G) Western blots of the gradient fractions in panel E. The position of phosphorylated Ltv1 is indicated with the asterisk. (H) Quantification of the Northern blots in panel F and a biological replicate. (I) Quantification of the Western blots in panel G and a biological replicate. Data are shown as a mean with standard deviation. (J) Normalized (to wt Rio2) doubling times from yeast expressing wt Ltv1 or Ltv1_YDY, and wt Rio2, or Rio2_K105E, or Rio2_D253A. Significance was tested using a two-way ANOVA. **, P<0.01; ****, P<0.0001.

To test this hypothesis, we took advantage of the observation that the Ltv1 phosphosite is mispositioned in Ltv1_L216S [9]. We therefore tested for genetic interactions between Ltv1_L216S and Ltv1_YDY. The data in **Figure 5C** show that these two mutations are epistatic, indicating that perturbing the location of the phosphosite (via the Ltv1_L216S mutation) perturbs its interaction with l31 (which is perturbed by Ltv1_YDY).

To further confirm that Ltv1 itself relays information between its phosphorylation status and Rio2 via interaction with l31, we combined the Ltv1_YDY mutation, which alters the interactions at l31 with the Rps20_DE and Rps3_KK mutations. Indeed, the Ltv1_YDY mutation fully rescues the detrimental growth defects from the Rps3_KK mutation and largely rescues the Rps20_DE mutation (**Figure 5D)**.

Next, we wanted to test if the Ltv1_YDY mutation also rescued the phosphorylation and release defects of Ltv1 that we have previously observed with Rps3_KK [20]. We therefore used sucrose gradient analysis in Fap7 depleted yeast as described above (**Figure 5E-G**). Indeed, while Ltv1 is not phosphorylated in cells with Rps3_KK as we have previously seen, when combined with Ltv1_YDY, Ltv1 phosphorylation is rescued (**Figure 5G**). Thus, changes in l31 are communicated to the Ltv1 phosphorylation site.

Despite the apparent full rescue of the Ltv1 phosphorylation, Ltv1 release was not rescued in the Ltv1_YDY variant (**Figure 5G&I**). Consistently, both Enp1 and Rio2, whose release requires prior Ltv1 release also remain in pre-40S ribosomes, and the formation of 80S-like ribosomes is only partially rescued (**Figure 5F-H**). The observation that Ltv1 phosphorylation but not release is rescued by the Ltv1_YDY mutation demonstrates the importance of l31 (where YDY binds) for Ltv1 release *after* it has been phosphorylated, and strongly suggests that rearrangements in Ltv1 communicate its phosphorylation to l31.

We have recently shown that Ltv1 release requires Tsr1’s movement from h44 to the beak [9, 16]. Moreover, this movement integrates information on Rio2’s and Ltv1’s phosphorylation state [20]. To further test the importance of Ltv1’s interactions with l31 for receiving the input from Rio2 phosphorylation, we combined the Ltv1_YDY mutation with a mutation that inactivates the Rio2 kinase activity, Rio2_D253A, as well as with a mutation in Rio2 (Rio2_K105E) that blocks the movement of Tsr1 from the decoding helix to the beak [9, 16]. Notably, Ltv1_YDY partially rescued the growth defect from K105E and is epistatic with Rio2_D253A (**Figure 5I**). This observation provides strong support for the model that changes in the interaction between Ltv1 and l31 (by Ltv1_YDY) are required for the shift in Tsr1 that is required for Ltv1 release. Moreover, the data also suggest that Rio2 phosphorylation affects Ltv1 via l31.

### Rio2 phosphorylation is communicated to Tsr1 via Rps15

In addition to the interaction of both Rio2 and Ltv1 with l31, which is required for Tsr1 repositioning and Ltv1 release, the Tsr1 position is linked to Ltv1 via Rps15 [9, 16]. In particular, our previous data have provided evidence that the C-terminal tail (CTT) of Ltv1 interacts with residues Y123, R127 and R130 of Rps15 (Rps15_YRR, **Figure 6A**), while residues R137E and K142E (Rps15_RK) bind Tsr1 residues R709 and K712 (Tsr1_RK), which are also positioned adjacent to l31 (**Figure 6B**). This was supported by epistatic interactions between truncation of Ltv1to remove the CTT (Ltv1_394) and Rps15_YRR, as well as Tsr1_RK and Rps15_RK, as well as synthetically sick interactions between Tsr1_RK and Rps15_YRR and Rps15_RK and Ltv1_394 [9, 16]. To test if Rio2 was part of this network and therefore regulated Tsr1 repositioning in response to its phosphorylation, we combined deletion of the N-terminal domain of Rio2 (Rio2_ΔN), which is located adjacent to Rps15, with Rps15_YKK and Rps15_RK. While Rio2_ΔN rescues the Rps15_YRR mutation, it does not have a substantial genetic interaction with Rps15_RK (**Figure 6C**). In contrast, mutation of Rio2_K105E, which blocks the release of Ltv1 [16], because it blocks repositioning of Tsr1 [9], is synthetically sick with Rps15_YRR and again does not demonstrate genetic interactions with Rps15_RK (**Figure 6D**). Together, these data suggest that Rio2 phosphorylation is communicated to Tsr1 for its repositioning via Rps15.

**Figure 6:**
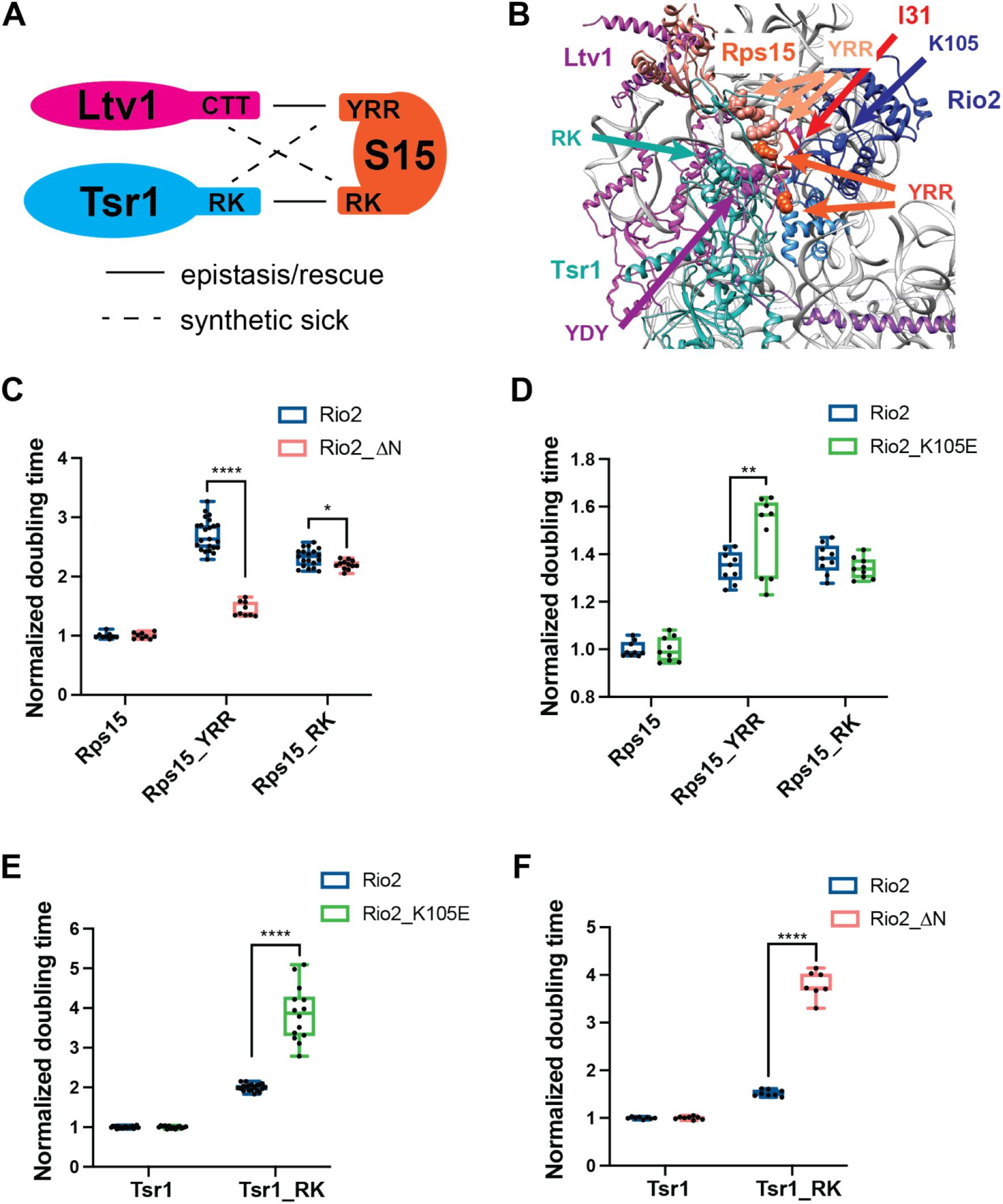
Rio2 is part of a genetic and structural network between Ltv1, Tsr1 and Rps15. (A) Genetic network between Rps15, Ltv1 and Tsr1 (adapted from [9]). (B) Detail of pre-40S ribosomes (pdb ID 6FAI), and Ltv1 placed as in [9]. Tsr1_RK is in cyan spacefill, Rio2_K105E is in blue spacefill, and Ltv1_YDY is in purple spacefill. Rps15_YRR and Rps15_RK residues are in salmon and orange, respectively. The position of l31, which is disordered in pre-40S is shown in red from mature 40S (PDB ID 3jam). (C) Normalized (to wt Rps15) doubling times from yeast expressing wt Rio2, or Rio2 lacking the N-terminal domain (aa 1-85, Rio2_ ΔN), and wt Rps15 or Rps15_YRR, or Rps15_RK. Significance was tested using a two-way ANOVA. **, P<0.01. (D) Normalized (to wt Rps15) doubling times from yeast expressing wt Rio2, or Rio2_K105E and wt Rps15 or Rps15_YRR, or Rps15_RK. Significance was tested using a two-way ANOVA. ****, P<0.0001. (E) Normalized (to wt Tsr1) doubling times from yeast expressing wt Rio2, or Rio2_K105E and wt Tsr1 or Tsr1_RK. Significance was tested using a two-way ANOVA. ****, P<0.0001. (F) Normalized (to wt Tsr1) doubling times from yeast expressing wt Rio2, or Rio2_ ΔN and wt Tsr1 or Tsr1_RK. Significance was tested using a two-way ANOVA. ****, P<0.0001.

Moreover, we note that Tsr1_RK and Rio2_K105E block repositioning of Tsr1 in different ways, as these are synthetically sick (**Figure 6E**). Similarly, Rio2_ΔN is sick with Tsr1_RK **(Figure 6F**). Together, these data strongly suggest that Rio2 phosphorylation and Ltv1 positioning are independent inputs into Rps15 to effect Tsr1 rotation.

## Discussion

### Binding of Asc1 initiates the late 40S assembly cascade by regulating Hrr25 phosphorylation

Previous work has revealed that entry into the cascade of events that quality control the assembly of the small subunit head is gated by Hrr25-dependent phosphorylation of the assembly factor Ltv1, which then converts a stable assembly intermediate into a series of unstable intermediates that ultimately complete maturation via an 80S-like assembly intermediate (**Figure 1C**, [9, 12–16]. What was unclear was how Hrr25-dependent phosphorylation of Ltv1was regulated.

Conversely, while published data from the Pertschy and our lab indicated that formation of a contact between Rps3 and Rps20 was necessary for this cascade [13], and in particular the Ltv1 phosphorylation step [16], genetic, biochemical and structural data had also indicated that in the stable intermediate, this contact was not yet made [9, 25], indicating that Hrr25-dependent phosphorylation of Ltv1 itself was regulated.

Here we show that by binding Rps3, Asc1 mediates repositioning of Ltv1 (and likely Rps3), to promote the formation of the contact between Rps3 and Rps20, thus enabling Ltv1 phosphorylation and committing the nascent subunits to the final maturation steps. Our data also indicate that Asc1 is aided in this by Rps17, which similarly interacts with Rps3. Analysis of the available structures and structure probing data suggests possible structural mechanisms for relaying the information about Asc1 binding to Ltv1 (**Figure 7A-C**): A cryo-EM structure from the Plisson-Chastang lab demonstrated that prior to dissociation of Ltv1, Rps3 is not in the location it adopts in mature 40S ribosomes [14], consistent with previous biochemical data [13]. While the disorder in the area of interest precludes the precise location of the protein, it is clear that in particular the C-terminal KH-domain of Rps3 is moved away from the ribosome (**Figure 7B**). Rps17 and Asc1 both bind the C-terminal tail that emanates from this KH-domain. In addition, there is no density for the N-terminal helix of Rps3, which in mature 40S makes a contact with Rps20, and which has been tentatively modeled in an alternative location. Consistently, we have previously shown that this interaction between Rps3 and Rps20 is not formed when Ltv1 is present [25], and a more recent placement of Ltv1 on pre-40S strongly suggests that Ltv1 actively blocks this interaction, by locating where this N-terminal helix in Rps3 will bind in mature 40S [9]. Thus, a plausible model could be that binding of Asc1 to Rps17 and the C-terminal tail of Rps3 stabilizes the Rps3 C-terminal tail via the tail’s interaction with both Rps17 and Asc1, and that this would drag the KH domain towards its native binding site (**Figure 7A, C)**. In that location, it would clash with a different portion of Ltv1 (**Figure 7C**), thus displacing Ltv1 in that area, allowing for the contact of Rps3 and Rps20 to be made.

**Figure 7:**
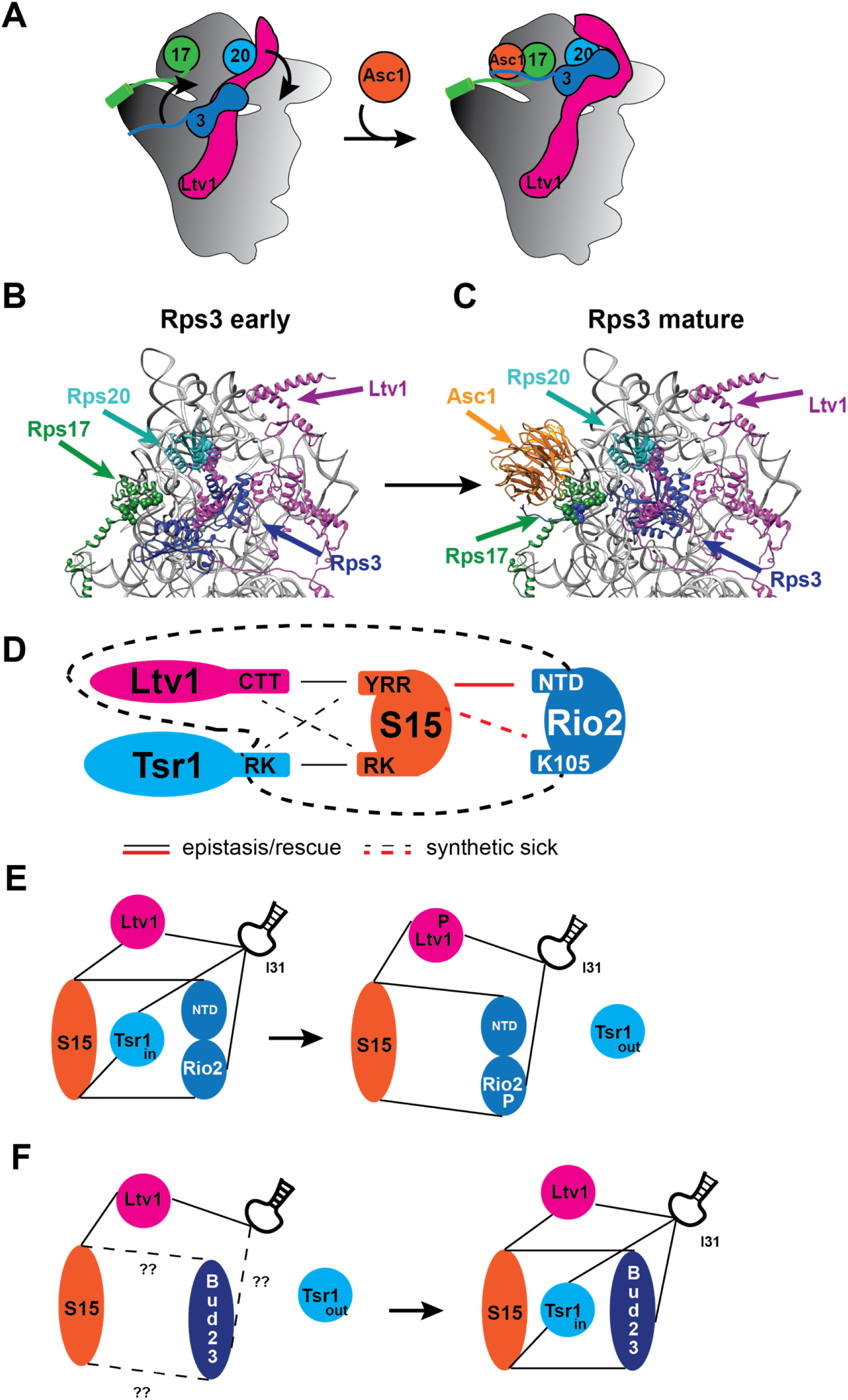
Ltv1 orchestrates the hierarchy in assembly of the 40S head. (A) Schematic illustration of the Rps17- and Asc1-mediated repositioning of Rps3 and Ltv1 to allow for the contact between Rps3 and Rps20, which enables Ltv1 phosphorylation. (B) Highlights of the structures of pre-40S ribosomes prior to the formation of the Rps3, Rps20 contact (accumulated with the Rps20_DE mutation). From PDB ID 6RBD. (C) Highlights of the structures of pre-40S ribosomes after the formation of the Rps3, Rps20 contact. From PDBID 3JAM. (D) Genetic network between Rio2, Rps15, Ltv1 and Tsr1. (E) Ltv1 and Rio2 phosphorylation are both communicated to Tsr1 via l31; both also communicate to Tsr1 via Rps15, creating a “fail-safe” network to regulate Tsr1 movement from h44 (in) to the beak (out). (F) Speculative analogous interactions between Bud23 and l31, and Rps15 might regulate the opposite movement of Tsr1 during binding.

### Tsr1 movement to the beak requires a dual input

We have previously shown that the formation of 80S-like ribosomes is associated with a rotation of Tsr1 from the decoding site helix towards the beak [9]. This appears to be the reverse movement it makes when binding [28]. In addition, we have also shown that this movement of Tsr1 is required to release Ltv1 from nascent subunits, and to allow for the binding of Rps31 [9]. Those data also demonstrated that Rps15 has a role in regulating this movement in response to changes in Ltv1 [9]. Here we expand this regulatory network and demonstrate that Rio2 also impacts the Tsr1 movement, again via Rps15 (**Figure 7D**). Thus, Rps15 emerges as a key regulator of Tsr1 movement, responding to dual inputs from both Ltv1 and Rio2. Moreover, our data also indicate that both Rio2 and Ltv1 bind loop 31 in 18S rRNA (l31), and that at least the interaction by Ltv1 is also required for Tsr1 movement and dissociation. Together, these observations explain how Ltv1 release requires the phosphorylation of both Ltv1 and Rio2. Moreover, this dual-fail safe mechanism also indicates the importance of the regulated Tsr1 movement, which enables both Ltv1 release and Rps31 binding, and ultimately the final maturation steps of the nascent 40S subunit.

The Tsr1 movement from h44 to the beak to enable Ltv1 dissociation appears to be the reverse of the movement by which it binds [28]. Tsr1 recruitment to nascent 40S occurs after Ltv1 binding and coincides with the binding of Rps15 [29, 30]. We thus speculate that the network of Ltv1 and Rps15 regulates the recruitment of Tsr1, akin to how it regulates its dissociation (**Figure 7E-F)**. Perhaps Bud23, which adopts the position of Rio2 in earlier pre-40S, could play the analogous role that Rio2 plays for Tsr1 release during its recruitment.

### Ltv1’s extended, flexible structure allows for the communication of assembly events across the nascent subunit

The assembly and quality control of the small subunit head involves a hierarchical cascade of events that is initiated by binding of Rps17 and Asc1, which enable the contact between Rps3 and Rps20, allowing for phosphorylation of Ltv1. The final step in this cascade is the formation of 80S-like ribosomes (**Figure 1C)**. While previous work has revealed the many steps involved, as well as the interactions that are necessary for each step [12–16, 20], what has remained unclear is how the successful completion of one event is communicated to allow the next step. In here we show that the assembly factor Ltv1 mediates the communication of each of the steps that occur while it remains bound to the subunit. We have previously shown that Ltv1’s extended, non-globular and flexible structure extends across the small subunit head, contacting key areas that are important for this communication, including Rps3/Rps20 near their contact point, j34 and l31, in whose folding it is involved in [9]. Here we show using a combination of genetic and biochemical data that Ltv1 itself communicates each of the events in this cascade (formation of the Rps3/Rps20 contact, phosphorylation of Ltv1, phosphorylation of Rio2). This is likely enabled not only by the extended conformation of Ltv1, but also the structural plasticity that arises because of its non-globular structure, which could allow it to adopt multiple binding poses on the subunit.

Notably, Ltv1 is not the only assembly factor that lacks a globular structure, and instead extends over the entire surface of the molecule. Utp14, Nop14, Mpp10 and Utp11 are 40S assembly factors with similar properties and Nog1, Nog2, and Erb1 are 60S assembly factors with large unstructured extensions. We speculate that these might similarly use their structural plasticity to create hierarchy in other stages of ribosome assembly. Interestingly, all of the above non-globular AFs are connected either by direct interactions, genetic data, or biochemical evidence to nucleotide-hydrolyzing enzymes. Ltv1 is regulated by Hrr25, and its phosphorylation regulates the Rio2 kinase activity. Utp14 regulates the DExH ATPase Dhr1, while Utp11, Nop14 and Mpp10 all bind the GTPase Bms1, indicating that this potential for conformational change is commonly exploited for regulatory purposes.

### Ltv1 mutants that disturb communication in head assembly are associated with disease

While its structural plasticity enables the hierarchy in the cascade that governs head assembly and its quality control, it also results in Ltv1’s vulnerability to mutations that disturb this careful balance of conformational shifts. The Ltv1_L216S mutation, which is associated with LIPHAK syndrome [31], repositions Ltv1 globally [9] and leads to global defects in head assembly. Moreover, the residues mutated in Ltv1_YDY, are also mutated in cancer patients with the mutation at the second Y recurring [32]. Additionally, mutations related to those mutated in Ltv1_FMS (F341A, M434A and S344A) are also mutated in human cancer cells. Human Ltv1_S244, which corresponds to S342 in yeast and is part of the phosphosite, is mutated recurrently. Moreover, E234, S238 and R239 are also mutated in humans, with recurring mutations in the arginine. These residues correspond to yeast K332, S336 and D337. A similar mutation in yeast, Ltv1_AMD (A334W, M335S, D337R, corresponding to human T236, K237 and R239), rescues the Rps3_KK and Rps20_DE mutants, analogously to the Ltv1_FMS mutant (**Figure S1**). Thus, the unique non-globular and flexible structure of Ltv1 not only allows for the facile communication and integration of multiple assembly steps, thereby establishing quality control checkpoints, it can also establish a vulnerability that renders cells sensitive to mutations that might bypass these steps. We speculate that these mutants might be prone to loss of Asc1/RACK1, as they can bypass the Asc1-motivated contact between Rps3 and Rps20.

## Materials and Methods

### Yeast Strains and Plasmids

*Saccharomyces cerevisiae* strains (**Table S1)** were either obtained from the Horizon Dharmacon Yeast Knockout Collection, or made by homologous PCR recombination [33] and confirmed by serial dilution, PCR, and Western blotting when antibodies were available. Plasmids were generated by standard cloning techniques and confirmed via Sanger sequencing. The plasmids used are listed in **Table S2**.

### In Vivo Subunit Joining Assay

Gal:Fap7 cells (with the appropriate additional genomic alterations) were grown in YPD for >16h at 30°C to deplete Fap7, harvested in the presence of cycloheximide [16, 17, 19], and lysed under liquid N_2_. Cell debris were spun out and the supernatant was loaded onto a sucrose gradient as described below.

### Sucrose-Gradient Fractionation

5,000-7,500 OD of clarified lysate were loaded onto a 10-50% sucrose-gradient before centrifuged at 40,000 rpm for 2 hours using a SW41Ti rotor. The gradients were then fractionated into 700 µl fractions. The fractions were later used for Northern Blotting and Western Blotting. Northern blot samples were prepared by phenol chloroform extraction on 200 µl of each fraction. Western blotting samples were prepared by mixing each individual fraction with SDS loading buffer, ran on an SDS-PAGE gel, prior to transferring and probing the membrane with the appropriate antibodies, as explained in [16, 17, 19].

### Northern Blot

Northern blots were prepared from the gradient fractions, RNA was isolated through phenol/chloroform RNA extraction, denatured in formaldehyde, and separated on an agarose gel. The following probes were used:

20S: GCTCTCATGCTCTTGCC;
18S: CATGGCTTAATCTTTGAGAC;
25S: GCCCGTTCCCTTGGCTGTG.

### Growth Curve Measurements

Cells were grown in glucose minimal media or YPD overnight, before being diluted into fresh minimal media or YPD for 3-6 hours. The cultures were then diluted into a 96-well plate at a starting OD of 0.04-0.1 (depending on growth rate) and placed in Synergy.2 plate reader (BioTek) to record OD_600_ for 24-48 hours at 30°C, while the plate was shaking. To extract doubling times, reads between OD 0.08-0.18 were fit with single exponential curves using Prism.

### Antibodies

Antibodies for assembly factors were raised in rabbits by Josman LLC against purified recombinant protein. The antibodies were tested against recombinant protein and yeast cell lysate.

## Acknowledgements

This work was supported by National Institute of Health grant R35-GM136323 and HHMI Faculty Scholar Grant 55108536 to K.K. and F32-GM139302 to Y.Y. The funders had no role in study design, data collection and analysis, decision to publish, or preparation of the manuscript. We thank members of the Karbstein lab for discussion and comments on the manuscript.

---

**Table S1:** Yeast strains used in this study.

| Strains | Description | Genotype | Reference |
| --- | --- | --- | --- |
| YKK73 | ΔLtv1 | BY4741 (MATa His3-1 Leu2-0 Met15-0 Ura3-0), Ltv1::KAN | [19] |
| YKK493 | Gal:S3 | BY4741, Gal:Rps3 (KAN) | [16] |
| YKK1275 | Gal:S3, Gal:Fap7 | BY4741, Gal:Rps3 (NAT), Gal:Fap7(KAN) | [16] |
| YKK422 | ΔLtv1, Gal:Fap7 | BY4741, Ltv1::KAN, Gal:Fap7(NAT) | [19] |
| YKK1502 | ΔAsc1, Gal:Fap7, | BY4741, Asc1::KAN; Gal:Fap7(Hyg) | This work |
| YKK729 | ΔLtv1, Gal:S20 | BY4741, Ltv1::KAN, Gal:Rps20 (HYG) | [17] |
| YKK762 | ΔLtv1, Gal:S3 | BY4741, Ltv1::KAN, Gal:Rps3 (NAT) | [36] |
| YKK500 | ΔAsc1, Gal:S3 | BY4741, Asc1::KAN, Gal:Rps3 (NAT) | This work |
| YKK1305 | ΔLtv1, Gal:S3, Gal:Fap7 | BY4741, Ltv1::hyg, Gal:Rps3 (NAT); Gal::Fap7 (KAN) | [17] |
| YKK723 | Gal:S3; Gal:S20 | Gal:Rps3 (KAN); Gal:Rps20 (HYG) | This work |
| YKK724 | Gal:S17; Gal:Fap7 | BY4741, ΔRps17A::KAN ; Gal:Rps17B(NAT), Gal:Fap7 (Hyg) | This work |
| YKK802 | ΔLtv1, Gal:S17 | BY4741, Ltv1::NAT, ΔRps17A::KAN ; Gal:Rps17B (Hyg) | This work |
| YKK1117 | ΔLtv1, Gal:S15 | BY4741, Ltv1::KAN, Gal:Rps15 (NAT) | [9] |
| YKK1190 | ΔLtv1, Gal:Rio2 | BY4741, Gal:Rio2(KAN), Ltv1::HYG | [9] |
| YKK1543 | ΔLtv1, ΔAsc1 | BY4741, Ltv1::Hyg, Asc1::Kan | This work |
| YKK642 | Gal:Rio2, Gal:Rps15 | BY4741, Gal:Rio2(KAN), Gal:Rps15 (NAT) | [9] |
| YKK1184 | Gal:Rio2, Gal:Tsr1 | BY4741, Gal:Rio2(KAN), Gal:Tsr1(NAT) | [16] |

**Table S2:** Plasmids used in this study.

| Plasmid | Description | Vector | Reference | Detailed description |
| --- | --- | --- | --- | --- |
| PKK3350 | wt Rio2 | TEF 413 | [16] |  |
| PKK3795 | Rio2_K105E | TEF 413 | [16] | Blocks Tsr1 movement |
| PKK3453 | Rio2_ΔN | TEF 413 | [16] | N-terminal deletion of aa 1-86 |
| PKK3795 | Rio2_D253A | TEF 413 | [16] | Phosphorylation-deficient |
| PKK3295 | wt Tsr1 | TEF 416 | [27] |  |
| PKK3716 | Tsr1_RK | TEF 416 | [16] | R709E,K712E, blocks Tsr1 movement |
| PKK30183 | wt Tsr1 | TEF 415 | [16] |  |
| PKK30184 | Tsr1_RK | TEF 415 | [16] | R709E,K712E, blocks Tsr1 movement |
| PKK30116 | Wt Rps15 | TEF 416 | [16] |  |
| PKK30180 | Rps15_YRR | TEF 416 | [16] | Y123I,R127K,R130K |
| PKK30181 | Rps15_RK | TEF 416 | [16] | R137E,K142E |
| PKK3890 | wt Rps20 | TEF 415 | [16] |  |
| PKK3934 | Rps20_DE | TEF 415 | [9] | D113A,E115A |
| PKK3891 | Rps20_EYER | TEF 415 | [16] | E80K,Y82A,E83K,R85E |
| PKK3606 | wt Ltv1 | TEF 413 | [17] |  |
| PKK30749 | Ltv1_L216S | TEF 413 | [9] | L216S |
| PKK3693 | Ltv1_1-394 | TEF 413 | [16] | Deletion of residues after 394. |
| PKK30523 | Ltv1_YDY | TEF 413 | [9] | Y82A,D83R,Y84A |
| PKK30524 | Ltv1_FMR | TEF 413 | This work | F341A, M343A, S344A |
| PKK30327 | Ltv1_AMD | TEF 413 | This work | A334W,M335A,D336R |
| PKK3521 | wt Rps3 | TEF 416 | [17] |  |
| PKK3867 | Rps3_KK | TEF 416 | [9] | K7A,K10A |
| PKK3933 | Rps3_KK | TEF 416 | [9] | K7A,K10D |
| PKK3525 | Rps3_214 | TEF 416 | [25] | Truncation after aa 214 |
| PKK31217 | Rps3_214_KK | TEF 416 | This work | K7A,K10A, truncation after aa 214 |
| PKK31203 | Rps3_IEP | TEF 416 | [25] | I208A, E210R and P211A |
| PKK3522 | U24 | TEF 413 | [25] | U24 snoRNA |
| PKK3582 | Wt Rps17 | TEF 413 | [25] |  |
| PKK3775 | Rps17_LRYTQ | TEF413 | [25] | L16A,R19E,Y20A,T39A,Q42A |

**Figure S1:**
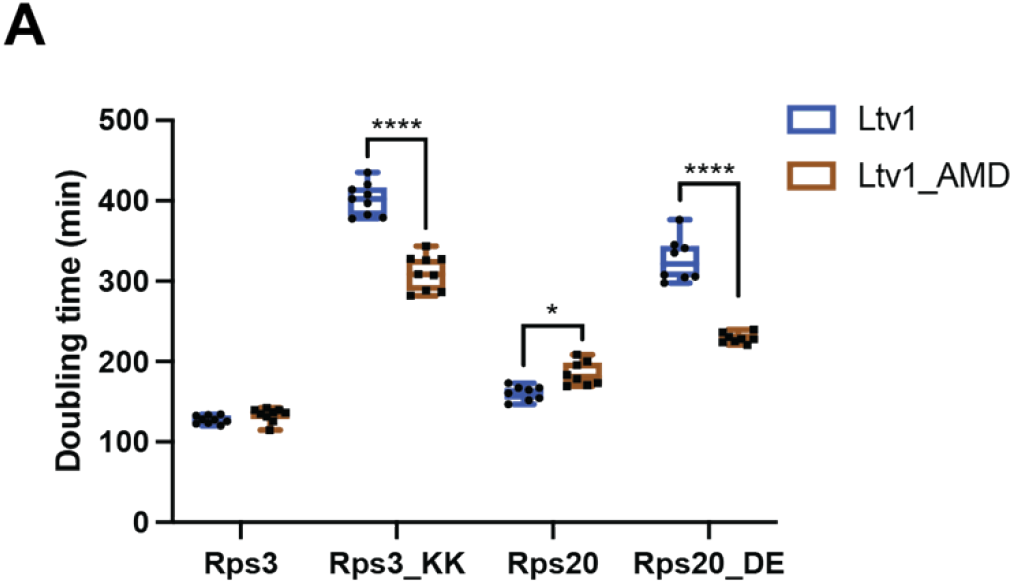
The cancer-associated Ltv1_AMD mutation allows for bypass of mutations that impair Ltv1 phosphorylation. Residues T236, K237 and R239 in human Ltv1 are mutated in cancer cells [32] and modeled by the yeast Ltv1_AMD mutation (A334W, M335S, D337R). Ltv1 is adjacent to Ltv1_FMS (F341A, M343A and S344A). Doubling times from yeast expressing wt Ltv1 or Ltv1_AMD and either wt Rps3, or Rps3_KK or wt Rps20 or Rps20_DE. Significance was tested using a two-way ANOVA. *, p<0.05, ****, P<0.0001.

